# A null model for Pearson coexpression networks

**DOI:** 10.1101/001065

**Authors:** Andrea Gobbi, Giuseppe Jurman

## Abstract

Gene coexpression networks inferred by correlation from high-throughput profiling such as microarray data represent a simple but effective technique for discovering and interpreting linear gene relationships. In the last years several approach have been proposed to tackle the problem of deciding when the resulting correlation values are statistically significant. This is mostly crucial when the number of samples is small, yielding a non negligible chance that even high correlation values are due to random effects. Here we introduce a novel hard thresholding solution based on the assumption that a coexpression network inferred by randomly generated data is expected to be empty. The theoretical derivation of the new bound by geometrical methods is shown together with applications in onco- and neurogenomics.

## 1 Introduction

Universally acknowledged by the scientific community as the basic task of the systems biology, the network inference is the prototypal procedure for moving from the classical reductionist approach to the novel paradigm of data-driven complex systems in the interpretation of biological processes [1]. The core of all the network inference (or network reconstruction) procedures is the detection of the topology of a graphy, *i.e.*, its wiring diagram, whose nodes are a given set of biological entities, starting from measurements of the entities themselves. In the last 15 years, the reconstruction of the regulation mechanism of a gene network and of the interactions among proteins from high-throughput data such as expression microarray of, more recently, from Next Generation Sequencing data has become a major line of research for laboratories worldwide. The proposed solutions rely on techniques ranging from deterministic to stochastic, and their number is constantly growing in the literature. Nonetheless, network inference is still considered an open, unsolved problem [2]. In fact, in many practical cases, the performances of the reconstruction algorithms are poor, due to several factors limiting the inference accuracy [3, 4] to the point of making it no better than coin tossing in some situations [5]. The major problem is the underdeterminacy of the task [6], due to the overwhelming number of interactions to predict starting from a usually small number of available measurements. In general, size and quality of available data are critial factors for all inference algorithms.

In what follows the impact of data size is discussed for one of the simplest inference techniques, *i.e.*, the gene coexpression network, where interaction strenght between two genes is a function of the correlation between the corresponding expression levels across the vailable tissue samples. The biological underlying hypothesis is that functionally related genes have similar expression patterns [7], and thus that coexpression is correlated with functional relationships, although this does not imply causality. In particular, as highlighted in [8], correlation can help unveiling the underlying cellular processes, since coordinated coexpression of genes encode interacting proteins, and Pearson correlation coefficient can be used as the standard measure. However, as noted in [9], correlation between genes may sometimes be due to unobserved factors affecting expression levels. Coexpression analysis has been intensively used as an effective algorithm to explore the system-level functionality of genes, sometimes outperforming much more refined approaches [10, 11]. The observation that simpler approaches such as correlation can be superior even on synthetic data has been explained by some authors [12, 13] with the difficulties of complex algorithm in detecting the subtleties of the combinatorial regulation. Moreover, coexpression network can capture more important features that the conventional differential expression approach [14], and its use has been extended to other tasks, for instance the investigation of complex biological traits [15]) Finally, these network can be crucial for understanding regulatory mechanisms [16], for the development of personalized medicine [17] or, more recently, in metagenomics [18].

Despite its success, a major issue affects coexpression networks: deciding when a given correlation value between two nodes can be deemed statistically significant and thus worthwhile assigning a link connecting them. This translates mathematically into choosing (a function of) a suitable threshold, as in the case of mutual information and relevance networks [19]. As reported in [20], in literature statistical methods for testing the correlations are underdeveloped, and thresholding is often overlooked even in important studies [21]. The two main approaches known in literature can be classified as soft or hard thresholding. The soft thresholding is adopted in a well-known framework called Weighted Gene Coexpression Network Analysis (WGCNA) [22], recently used also for other network types [23, 24]. All genes are mutually connected, and the weight of the link is a positive power of the absolute value of the Pearson correlation, where the exponent is chosen as the best fit of the resulting network according to a scale-free model [25, 26]. This approach, without discarding any correlation, promotes high correlation values and penalizes low values. In the hard thresholding approach, instead, only correlation values larger than the threshold are taken into account, and an unweighted link is set for each of these values, so that a binary network is generated (see [27] for one of the earliest references). Clearly, an uncorrectly chosen threshold value can jeopardize the discussed results with false negative links (for too strict threshold) or false positive links (for too loose threshold). Many different heuristics have been proposed for setting the threshold values, such as using the False Discovery Rate [28, 29, 30, 31], or the *p*-value of the correlation test [17], or employing partial correlation [32], or using rank-based techniques [33, 34, 35] or more complex randomization techniques [36]. Alternatively, correlation distribution has been studied, experimentally [37] or at level of single interaction, not as whole network [38]. However, in many studies in literature, the threshold is not chosen accordingly to a soundly bases procedure, but referring to standard choices [39, 40, 41, 42], or to heuristics not directly related to the correlation values, but rather with the resultining network topology [43, 44, 45, 46, 47, 48, 49, 50, 51]. In [52] a comparison of some coexpression thresholds is shown on a few microarray datasets.

Here we propose a new a priori and non-parametric model for the computation of an hard theshold based on the assumption that a random coexpression graph should not have any edge. The threshold is theorethically derived by means of a geometric approach based on the work of Bevington [53], and, as a deterministic independent null model, it depends only on the dimensions of the starting data matrix, with assumptions on the skewness of the data distribution compatible with the structure of gene expression levels data [54, 55]. By definition, this threshold is aimed at minimizing the possible false positive links, paying a price in terms of false negative detected edges.

To conclude with, we show four applications, in both the large and the small sample size settings. The first two are examples in a large sample size settings, with a synthetic dataset and with an ovarian epithelial carcinoma dataset on a large cohort of 285 cases [56, 57]. Two more applications in the opposite situations are demonstrated on two publicly available datasets, the former regarding a pancreatic cancer study [58] on a tiny cohort of six patients, and the latter on a Alzheimer dataset with 28 samples on two different phenotypes [59, 60, 61].

## 2 Distribution of Pearson correlation

Let *x, y* ∈ **R***^n^* with *n ≥* 3. The **Pearson correlation coefficient** *ρ* between *x* and *y* is defined as:

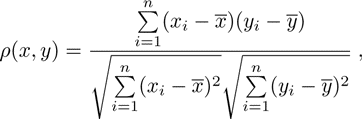

where *w̄* denotes the arithmetic mean 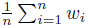 of the *n*-dimensional vector *w*.

The first step towards the construction of a null model for random absolute Pearson coexpression network is the estimation, for 0 *< p <* 1, of the function *F* (*n, p*) = *P* (|*ρ*(*x, y*)| *> p*), where *x* and *y* are two independent normal vectors of length *n*. Define two new random variable *x̃* and *ỹ* as follows:

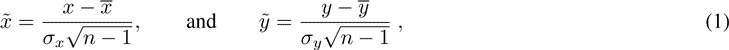

where *σ_x_* and *σ_y_* are the standard deviations of *x* and *y*. From the definition, the following identities immediately descend:

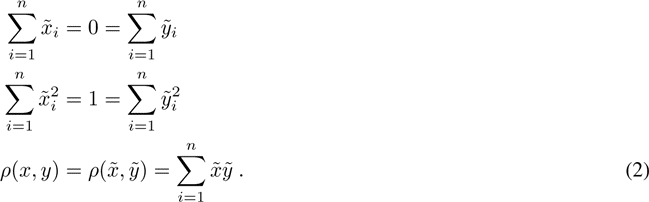

We can now state and prove two key results.

### Proposition 1.

*Let x, y, x̃*, *ỹ as in Eq. 1. Then x̃*, *ỹ* ∈ *S*_*n*−1_ ∩ *ℋ* ∼ *S*_*n*−2_, *where ℋ is the vectorial hyperplane defined as* 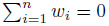 *and w_i_ are the coordinates of* **R**^*n*^.

*Proof.* Since 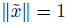, the following identity holds:

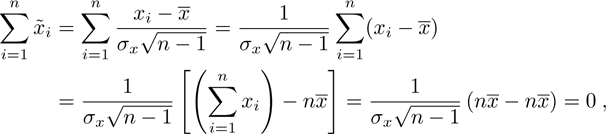

and the same holds for *ỹ*, too.

An example for *n* = 3 of the situation described in Prop. 1 is plotted in Fig. 2.

### Proposition 2.

*Let x, y as in Prop. 1 and 0 < p < 1 be a real number. Then the function F* (*n, p*) *has the following close form*

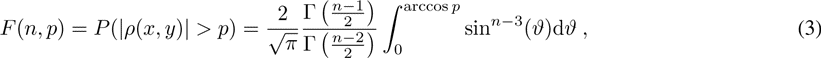

*where* Γ(*x*) *is the gamma function* 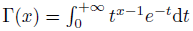

*Proof.* Using Eq. 1 and Eq. 2, we have that

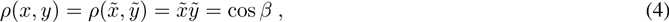

where *β* is the angle between the two vectors *x̃* and *ỹ*. Eq. 4 and Prop. 1 yields that *P* (*|ρ*(*x, y*)*| > p*) is the proportion between the area of the spherical cap in *n −* 2 dimensions included within an angle *β* from *x* and the whole surface of the *n −* 2-dimensional sphere [62]. A compact formula for the area 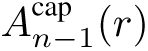 of a *n −* 2-spherical cap is given in [63] as:

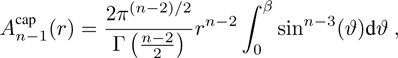

and, since the area of the whole surface is

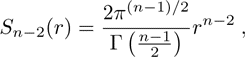

the thesis follows from the setting *r* = 1.

An alternative derivation of the same result can be found in [53].

In Prop. 1 the transformed vectors are assumed to be uniformly distributed on the spherical surface. This assumption holds in the case of a normal distribution, but it does not hold in general. However, in the following paragraph we sshow that is a good approximation, since *x* and *y* are independent. In fact, Prop. 3 can be generalised to other distributions [64, 65, 66, 67]), when data skewness can be bounded [62].

Let *G^δ^*(*p, n*) be an empirical distribution generated by *k* couples of two vectors *x, y ∈* **R***^n^* sampled according to a given distribution function *δ*. Let then

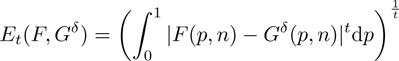

be the *t*-error function evaluating the difference between the theoretical distribution *F* (*p, n*) and the empirical distribution *G^δ^*(*p, n*). Hereafter we report the results of the simulations for *k* = 50000 and *n* = 8, 20, 100, where *δ* is one of the following three distribution functions:

- *U* (0, 1), the uniform distribution in [0, 1];
- *N* (*m, s*), the normal distribution with mean *m* and standard deviation *s*;
- *L*(*ml, sl*), the lognormal distribution with mean-log *ml* and standard deviation-log *sl*.

In particular, in Tab. 1 we list the values of *E*_2_(*F, G^δ^*) and in Fig. 1 we display the curves of the cumulative distribution functions (CDF) of *G^δ^*(*p, n*) corresponding to the three functions *δ*, separately for the different values of *n*.

**Figure 1:**
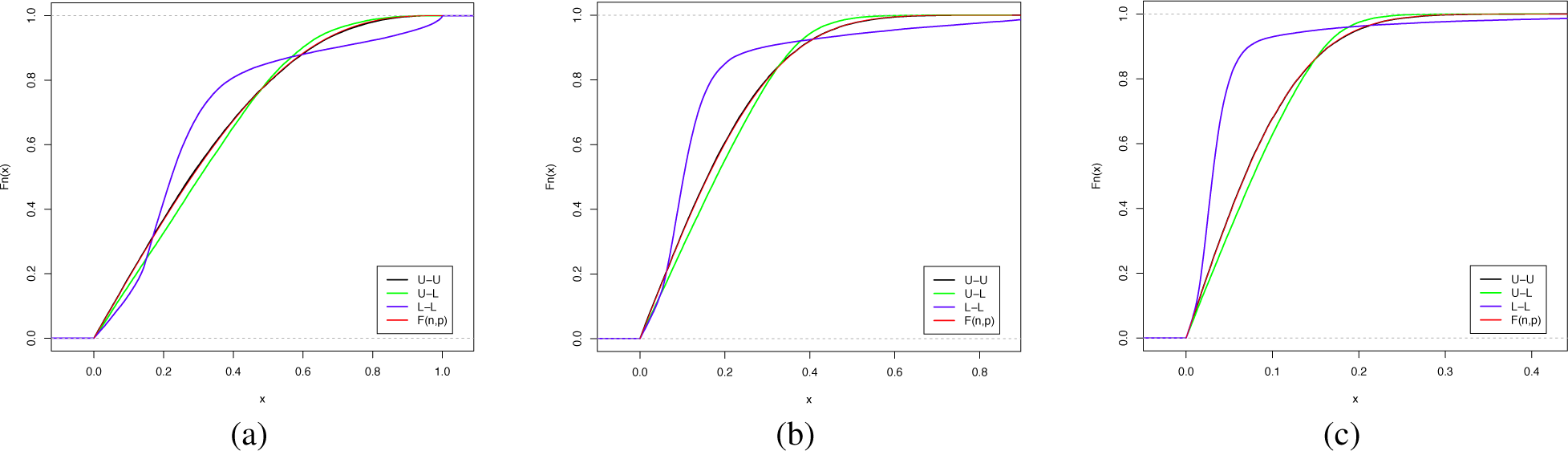
CDFs relative to the different distributions *δ* = *U* and *δ* = *L* compared with the theoretical curve *F* (*n, p*), for the three cases *n* = 8 (a), *n* = 20 (b) and *n* = 100 (c). In all cases, the red curve of *F* (*n, p*) and the black curve for the double uniform distribution *U − U* are almost coincident.

**Table 1:** Error function *E*_2_(*F, G^δ^*), for *n* = 8, 20, 100 and different distributions *δ*.

| $G^\delta(p, 8)$ | | $x$ | | |
| --- | --- | --- | --- | --- |
| | | $U(0, 1)$ | $N(0, 1)$ | $L(2, 3)$ |
| $y$ | $U(0, 1)$ | 0.001832 | 0.00137 | 0.021202 |
| | $N(0, 1)$ | 0.001195 | 0.00142 | 0.001432 |
| | $L(2, 3)$ | 0.022961 | 0.00139 | 0.080803 |
| $G^\delta(p, 20)$ | | $x$ | | |
| | | $U(0, 1)$ | $N(0, 1)$ | $L(2, 3)$ |
| $y$ | $U(0, 1)$ | 0.0016851 | 0.0007752 | 0.0248819 |
| | $N(0, 1)$ | 0.0008008 | 0.0014559 | 0.0008381 |
| | $L(2, 3)$ | 0.0238804 | 0.0011422 | 0.1038271 |
| $G^\delta(p, 100)$ | | $x$ | | |
| | | $U(0, 1)$ | $N(0, 1)$ | $L(2, 3)$ |
| $y$ | $U(0, 1)$ | 0.0006978 | 0.0008244 | 0.015630 |
| | $N(0, 1)$ | 0.0009281 | 0.0007388 | 0.001441 |
| | $L(2, 3)$ | 0.0159969 | 0.0014090 | 0.104998 |

Regardless of the value of *n*, the empirical distribution fits the exact formula Eq. 3 when *x* and *y* are uniformly sampled, while it does not fit the same equation when the two vectors come from extremely skewed distributions such as the lognormal. Note that non-Gaussian asymmetric distributions can occasionaly being detected in some array studies [55]: however, techinques for reducing the skewness are routinely applied during preprocessing [54], and thus the aforementioned results can be safely used in the microarray framework.

Finally, we conclude this paragraph deriving the mean and the variance of the function |*ρ|*. Starting from Eq. 3, the density function *f* (*p, n*) can be computed as

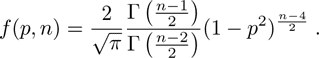

Using the above expression for *f* (*p, n*), the two moments follow straightforwardly:

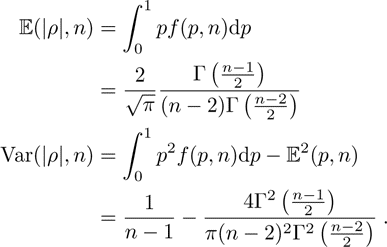

## 3 Coexpression network and threshold selection

The results derived in the previous section are used here to construct a null model for the correlation network, thus yielding a threshold for the inference of a coexpression network from nodes’ data.

Let χ = {_*i*_*x*}_*i*=1_^*m*^ be a set such that _*i*_*x* ∈ *𝒰*[0, 1]^*n*^ ∀*i* = 1,… *m*. Then the coexpression *p*-graph *𝒢*_*p*_ = {*V*, *E*_*p*_} is the graph where

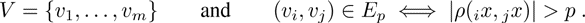

The first result characterizes the coexpression graphs in terms of null models:

### Proposition 3.

*The graph 𝒢*_*p*_ *is an Erdös-Rényi model [68] with m nodes and probability p as in Eq. 3*.

The proof follows immediately from the definition of *𝒢*_*p*_ and Eq. 3.

**Example** Consider a dataset *𝒴* consisting of *n* = 3 samples described by *m* = 100 genes. Then *𝒴* can be represented by 100 points in [0, 1]^3^ ⊂ **R**^3^ as shown in Fig. 2(a). The new variables _*i*_*x̃* are built through a two-stages procedure applied to each gene. First the mean is subtracted, so the transformed dataset lies on the hyperplane *ℋ* described in Prop. 1 as displayed in Fig. 2(b,c). Finally. each gene is normalized to unitary variance, and the resulting dataset lies on *S*_*n*−1_ ∩ *ℋ* which is the circumference in Fig. 2(d). Using the results in the previous section, it is now possible to define, for *n* nodes measured on *m* samples, the secure threshold *p̄* as the minimum value of *p* such that the corresponding random coexpression network *𝒢*_*p̄*_ is on average an empty graph, that is

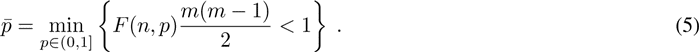

**Figure 2:**
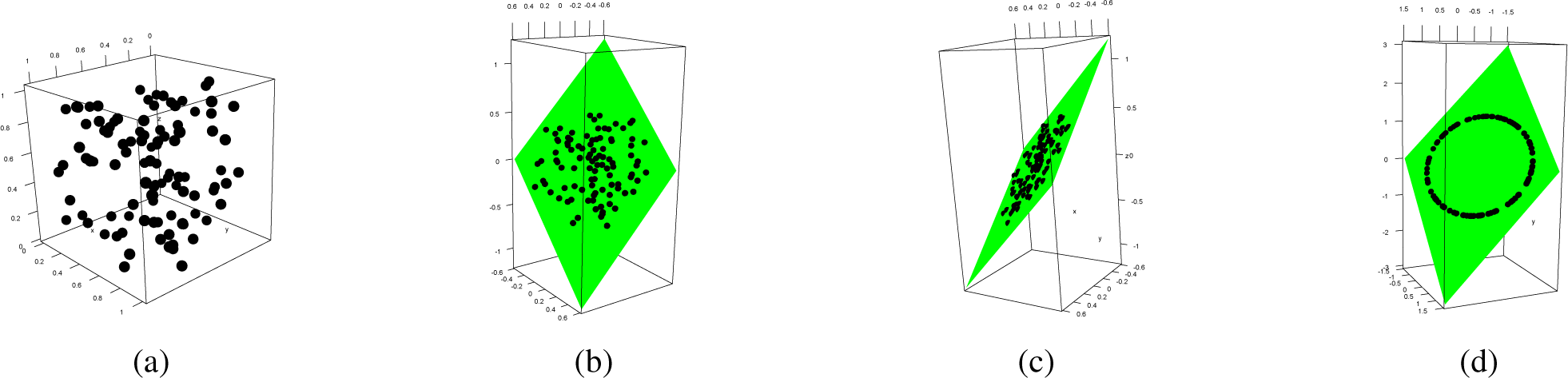
Transformation of the initial dataset preserving the Pearson correlation. (a) Original dataset (b,c) Mean substraction (d) Variance normalization. In green the hyperplane *ℋ*.

The underlying hypothesis for Eq. 5 is the assumption that in a random dataset we do not expect an kind of edge, *i.e.* any kind of relation. Due to its definition, the secure threshold *p̄* is biased towards avoiding the false positive links, paying a price in terms of false negatives. In fact, all the links passing the filter are induced by correlation only due to the inference data, while all links whose correlation value can be generated either by relation between data or by random noise are discarded. In Tab. 2 a collection of values of *p̄* is lisetd for different *m* and *n*, while in Fig. 3 the contourplot of the function *p̄*(*n*, *m*) is shown first on a large range of values and then zooming on the small sample size area. In the Tab. 3 we show the comparison on a set of synthetic and array datasets of the secure threshold *p̄* with another well known hard thresholding methods, the clustering coefficient-based threshold *C^∗^* [48] and with the statistical thresholds based on the adjusted p-values of 0.01, 0.05 or 0.1. In almost all cases, the threshold *p̄* is the strictest. As shown in the previous section, for not very skewed distribution, the good approximation provided by the exact formula for *F* (*n, p*) given in Eq. 3 guarantees the effectiveness of the secure threshold *p̄* in detecting actual links between nodes. Nonetheless, whenever a stricter threshold is needed, it is still possible to follow the construction proposed, with the following refinement. The edge-creation process in the Erdös-Rény model follows a binomial distribution, where *n* is the number of trials and *p* the probability associated to the succes of a trial. The mean *np* of this distribution is one of the contributing term in the definition of secure threshold Eq. 5. To further restrict the number of falsely detected links, the variance term (*np*(1 – *p*) for the binomial distribution) can be added to the formula through the Chebyshev’s inequality

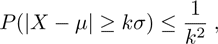

where *µ* and *σ* are the mean and the standard deviation of *X*. Thus, the definition of secure threshold can be sharpened to *p̃*_*k*_ as follows:

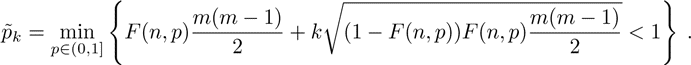

**Figure 3:**
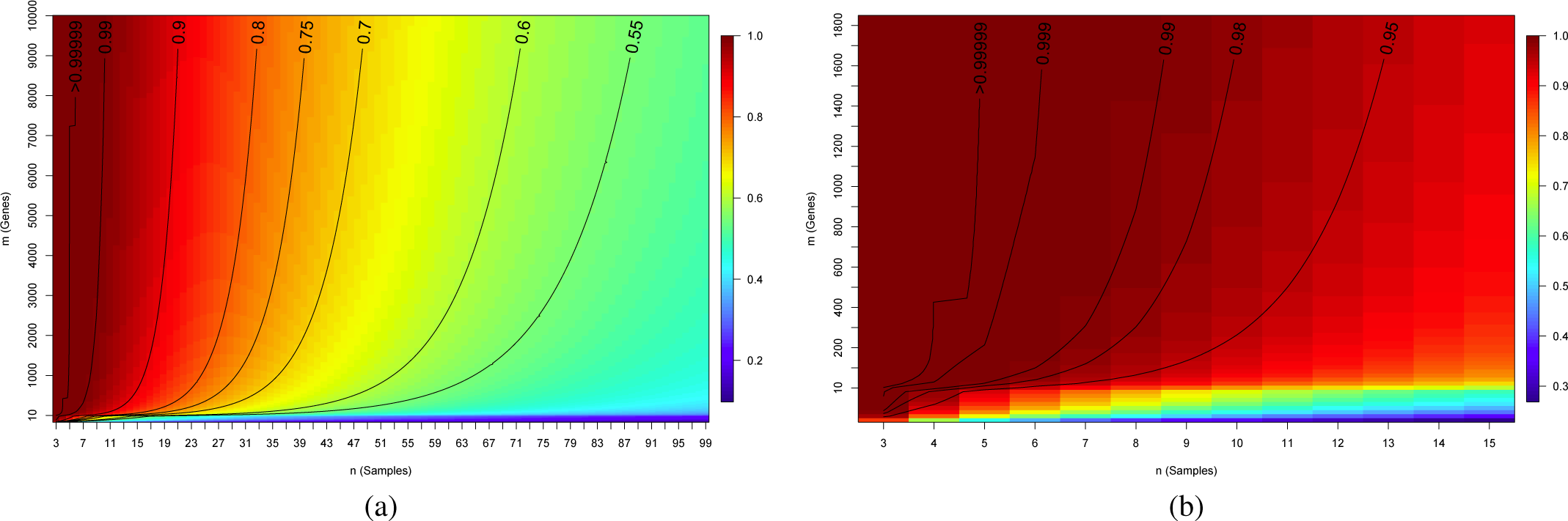
Contour plot of the function *p̄*(*m*, *n*) on (a) a large (*m, n*) range and (b) zoomed on the small sample size area.

**Table 2:** A subset of values of the secure threshold *p̄* for different number of samples *m* and genes *n*.

| $n \backslash m$ | 100 | 500 | 1000 | 2000 | 10000 | 50000 | 100000 |
| --- | --- | --- | --- | --- | --- | --- | --- |
| 8 | 0.95629 | 0.98520 | 0.99070 | 0.99415 | 0.99800 | 0.99932 | 0.99957 |
| 15 | 0.81681 | 0.89170 | 0.91323 | 0.93036 | 0.95800 | 0.97456 | 0.97949 |
| 20 | 0.73825 | 0.82388 | 0.85077 | 0.87330 | 0.91286 | 0.93973 | 0.94852 |
| 30 | 0.62814 | 0.71776 | 0.74817 | 0.77485 | 0.82534 | 0.86367 | 0.87729 |
| 50 | 0.50225 | 0.58534 | 0.61513 | 0.64213 | 0.69607 | 0.74036 | 0.75705 |
| 75 | 0.41647 | 0.49026 | 0.51740 | 0.54238 | 0.59353 | 0.63709 | 0.65394 |
| 100 | 0.36343 | 0.42999 | 0.45477 | 0.47774 | 0.52537 | 0.56662 | 0.58279 |

**Table 3:** Comparison of the secure threshold *p̄* with the clustering coefficient-based threshold *C^∗^* [48] and the statistical thresholds based on the adjusted p-values B0.01, B0.05 or B0.1 on a collection of synthetic and array datasets.

| Dataset type | #samples | #nodes | $C^*$ | B0.01 | B0.05 | B0.1 | $\bar{p}$ |
| --- | --- | --- | --- | --- | --- | --- | --- |
| Simulated | 50 | 1000 | 0.57 | 0.58 | 0.54 | 0.52 | 0.6152 |
| Simulated | 25 | 1000 | 0.69 | 0.76 | 0.72 | 0.70 | 0.7956 |
| H-U133P | 23 | 897 | 0.72 | 0.78 | 0.74 | 0.72 | 0.8125 |
| H-U133P | 10 | 897 | 0.78 | 0.96 | 0.94 | 0.93 | 0.9723 |
| H-U133P | 10 | 675 | 0.77 | 0.96 | 0.93 | 0.92 | 0.9681 |
| H-U133P | 9 | 897 | 0.79 | 0.97 | 0.96 | 0.95 | 0.9821 |
| H-U133P | 8 | 897 | 0.81 | 0.98 | 0.97 | 0.96 | 0.98999 |
| H-U133P | 7 | 897 | 0.81 | 0.99 | 0.99 | 0.98 | 0.99558 |
| H-U133P | 6 | 897 | 0.86 | >0.99 | >0.99 | 0.99 | 0.99872 |
| H-U133P | 5 | 897 | 0.92 | >0.99 | >0.99 | >0.99 | 0.99984 |
| H-U133P | 4 | 897 | 0.99 | >0.99 | >0.99 | >0.99 | > 0.99999 |
| H-U133A | 4 | 675 | 0.99 | >0.99 | >0.99 | >0.99 | > 0.99999 |
| H-I6 | 4 | 675 | 0.99 | >0.99 | >0.99 | >0.99 | > 0.99999 |
| M-U74 | 4 | 401 | 0.97 | >0.99 | >0.99 | >0.99 | 0.99999 |

For instance, the binomial distribution, for large value of *n*, can be approximated as a normal distribution for which the 95.45% of the values lie in the interval (*µ −* 2*σ, µ* + 2*σ*). In Tab 4 we show, for *p̃*_2_, the analogous of Tab. 2 for *p̄*. Finally, the Chebyshev’s inequality implies that at least the 96% of the values lie in the interval (*µ −* 5*σ, µ* + 5*σ*): the corresponding threshold values for *k* = 5 are listed in Tab. 5.

**Table 4:** A subset of values of the secure threshold *p̃*_2_ for different number of samples *m* and genes *n*.

| $n \backslash m$ | 100 | 500 | 1000 | 2000 | 10000 | 50000 | 100000 |
| --- | --- | --- | --- | --- | --- | --- | --- |
| 8 | 0.97584 | 0.99179 | 0.99484 | 0.99675 | 0.99889 | 0.99962 | 0.99977 |
| 15 | 0.86282 | 0.91826 | 0.93437 | 0.94723 | 0.96810 | 0.98065 | 0.98439 |
| 20 | 0.78966 | 0.85726 | 0.87876 | 0.89686 | 0.92883 | 0.95068 | 0.95784 |
| 30 | 0.68082 | 0.75573 | 0.78151 | 0.80425 | 0.84759 | 0.88074 | 0.89256 |
| 50 | 0.55034 | 0.62269 | 0.64902 | 0.67302 | 0.72135 | 0.76137 | 0.77651 |
| 75 | 0.45887 | 0.52436 | 0.54881 | 0.57145 | 0.61820 | 0.65834 | 0.67394 |
| 100 | 0.40153 | 0.46116 | 0.48369 | 0.50471 | 0.54865 | 0.58703 | 0.60214 |

**Table 5:** A subset of values of the secure threshold *p̃*_5_ for different number of samples *m* and genes *n*.

| $n \backslash m$ | 100 | 500 | 1000 | 2000 | 10000 | 50000 | 100000 |
| --- | --- | --- | --- | --- | --- | --- | --- |
| 8 | 0.98553 | 0.99508 | 0.99691 | 0.99805 | 0.99934 | 0.99978 | 0.99986 |
| 15 | 0.89287 | 0.93585 | 0.94842 | 0.95849 | 0.97486 | 0.98474 | 0.98768 |
| 20 | 0.82530 | 0.88080 | 0.89858 | 0.91361 | 0.94025 | 0.95853 | 0.96454 |
| 30 | 0.71934 | 0.78401 | 0.80647 | 0.82636 | 0.86445 | 0.89373 | 0.90420 |
| 50 | 0.58686 | 0.65162 | 0.67541 | 0.69720 | 0.74130 | 0.77803 | 0.79198 |
| 75 | 0.49164 | 0.55125 | 0.57373 | 0.59463 | 0.63803 | 0.67552 | 0.69015 |
| 100 | 0.43124 | 0.48595 | 0.50683 | 0.52640 | 0.56752 | 0.60368 | 0.61796 |

## 4 Applications in functional genomics

### 4.1 Large sample size

#### Synthetic dataset

First a correlation matrix *M*_*𝒢*_ on 20 genes *G*_1_, … *G*_20_ is created, together with a dataset *𝒢* of the corresponding expression *G*_*i*_^1000^ across 1000 synthetic samples, so that *M*_*𝒢*_(*i*, *k*) = |cor(*G*_*i*_^1000^, *G*_*j*_^1000^)| is the absolute Pearson correlation between the expression of the genes *G_i_* and *G_j_* from *𝒢*.

In particular, *M*_*𝒢*_ has two 10 × 10 blocks highly correlated on the main diagonal, and two 10 × 10 poorly correlated blocks on the minor diagonal, as shown in Fig. 4. These blocks derived from the following generating rule, given uncorrelated starting element *G*_1_^1000^ and *G*_11_^1000^:

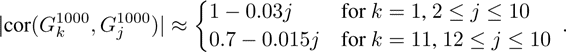

**Figure 4:**
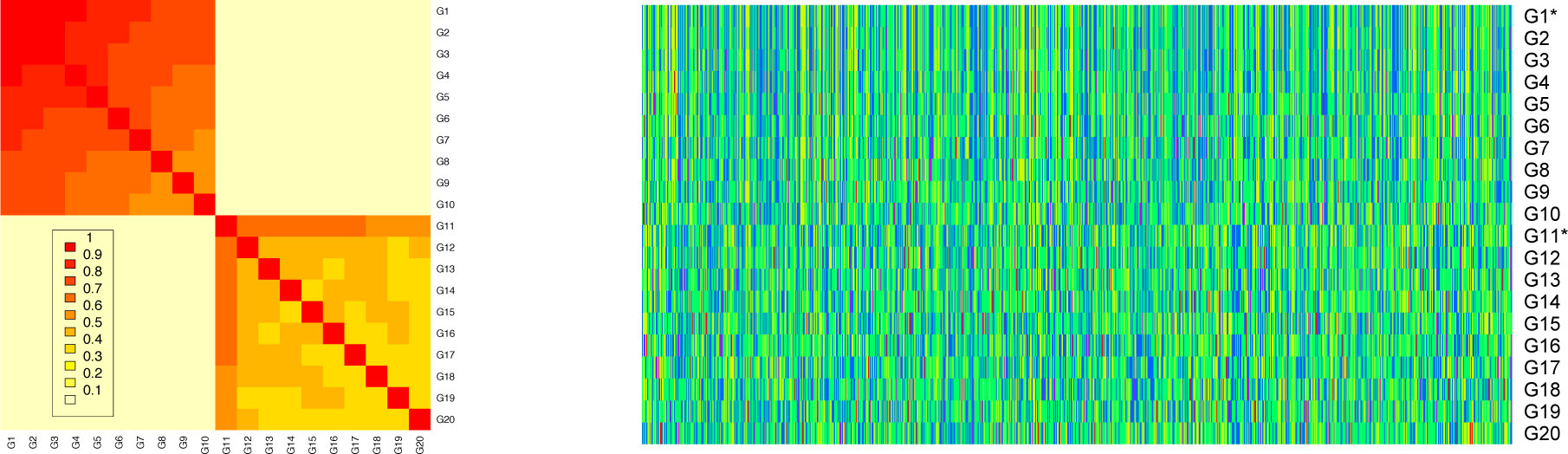
Levelplot of the structure of the correlation matrix *M*_*𝒢*_ (left) and heatmap of the dataset *𝒢*. The generating gene expression vectors *G*_1_^1000^ and *G*_11_^1000^ are marked with *.

Outside the two main blocks, all correlation values range between 0.002 and 0.074. In Fig. 4 we also show the heatmap of the gene expression dataset *𝒢*. Then a subset of *n_s_* samples is selected from the starting 1000, and the corresponding coexpression networks is built, for the 100 hard threshold values 0.01j, for 1 ≤ *j* ≤ 100. The secure threshold for these cases are respectively 0.799, 0.596 and 0.389. These procedure is repeated 500 times for each value *n_s_* = 10, 20, 50. The same experiment is then repeated adding a 20% and a 40% level of Gaussian noise to the original data. Using *M*_*𝒢*_ as the ground truth where all values outside the two main blocks are thresholded to 0, for each hard threshold 0.01j we evaluate the ratio of False Positive links, the ratio of False Negative links and the Hamming-Ipsen-Mikhailov (HIM) distance from the gold standard^1^ The graphs summarizing the experiments, separately for sample size, are displayed in Fig. 5.

**Figure 5:**
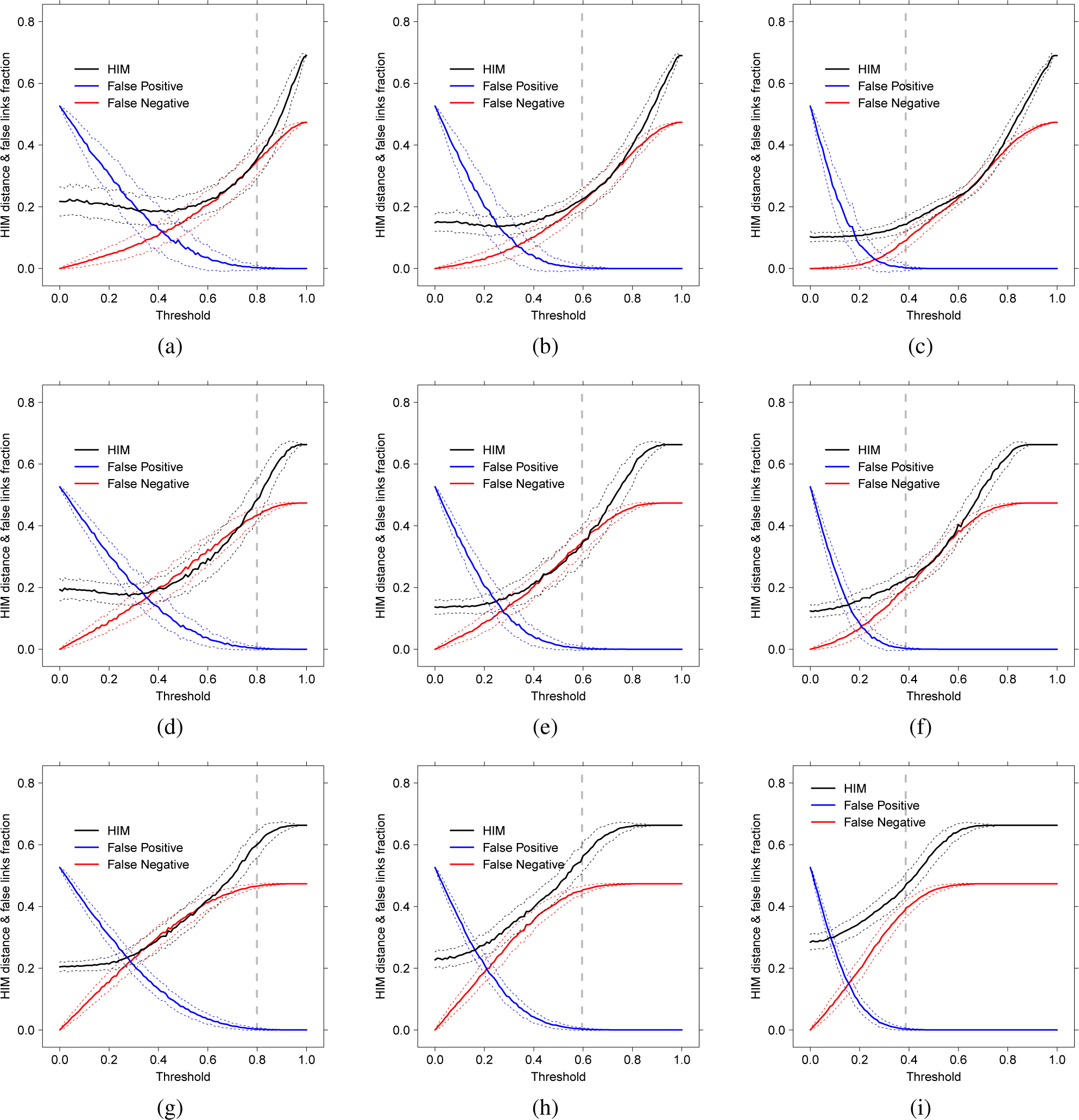
Coexpression inference of the *M*_*𝒢*_ network from random subsampling of the *𝒢* dataset, witouth noise (a,b,c), with 20% Gaussian noise (d,e,f) and with 40% Gaussian noise (g,h,i), on 10 (a,d,g), 20 (b,e,h) and 50 (c,f,i) samples. Solid lines indicate mean over 500 replicates of HIM distance (black), ratio of False Positive (blue) and ratio of False Negative (red); dotted lines of the same color indicate +/− 1*σ*, while grey vertical dashed lines correspond to the secure threshold *p̄*.

In all cases, the secure threshold *p̄* corresponds to the strictest value yielding a coexpression network with no false positive links included, which its characterizing property. Moreover, in almost all displayed situations, thresholding at *p̄* still guarantees an acceptable HIM distance from the ground truth, and a false negative ratio always smaller than 0.4.

#### Ovarian cancer

The aforementioned results obtained in a synthetic case are then tested here in a large array study on 285 patients of ovarian cancer at different stages [57], recently used in a comparative study on conservation of coexpressed modules across different pathologies [56]. In details, a whole tumor gene expression profiling was conducted on 285 predominately high-grade and advanced stage serous cancers of the ovary, fallopian tube, and peritoneum; the samples were hybridized on the Affymetrix Human Genome HG-U133 Plus 2.0 Array, including 54621 probes. The goal of the original study was to identify novel molecular subtypes of ovarian cancer by gene expression profiling with linkage to clinical and pathologic features. As a major result, the authors presented two ranked gene lists supporting their claim that molecular subtypes show distinct survival characteristics. The two gene lists characterize the Progression Free Survival (PSF) and the poor Overall Survival (OS), respectively.

Following the procedure of the previous, synthetic example, first we individuate the sample subset corresponding to the homogeneous cohort of 161 grade three patients and a set *T* of 20 genes, belonging to the top good OS and PFS genes (EDG7, LOC649242, SCGB1D2, CYP4B1, NQO1, MYCL1, PRSS21, MGC13057, PPP1R1B, KIAA1324, LOC646769) and to the top poor OS/PFS genes (THBS2, SFRP2, DPSG3, COL11A1, COL10A1, COL8A1, FAP, FABP4, POSTN), thus generating a dataset *𝒪*_*T*_ of dimension 161 samples and 20 features. The corresponding absolute Pearson correlation matrix *O_T_* is then used as the ground truth for the subsampling experiments: the levelplot of *O*_*T*_ and the heatmap of *𝒪*_*T*_ is shown in Fig. 6. In these experiments, a random subdataset of *n_s_* samples is extracted from *𝒪*_*T*_, and the corresponding absolute Pearson coexpression network on the nodes *T* is built, for increasing threshold values. In Fig. 7 we report the HIM and the ratio of False Positive and False Negative links for 500 runs of the experiments, separately for *n_s_* = 5, 10, 20 and 50.

**Figure 6:**
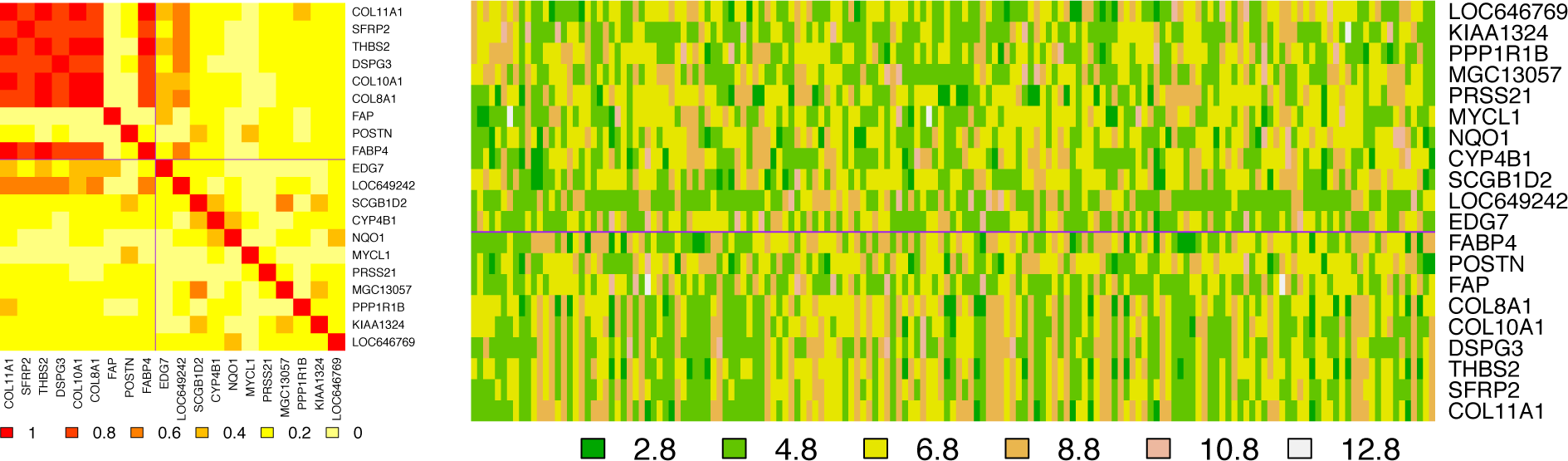
Levelplot of the structure of the correlation matrix *O_T_* (left) and heatmap of the Ovarian dataset *𝒪*_*T*_ restricted to the set of 20 selected genes *T*. Solid lines separate the group of good and poor PFS/OS top genes.

**Figure 7:**
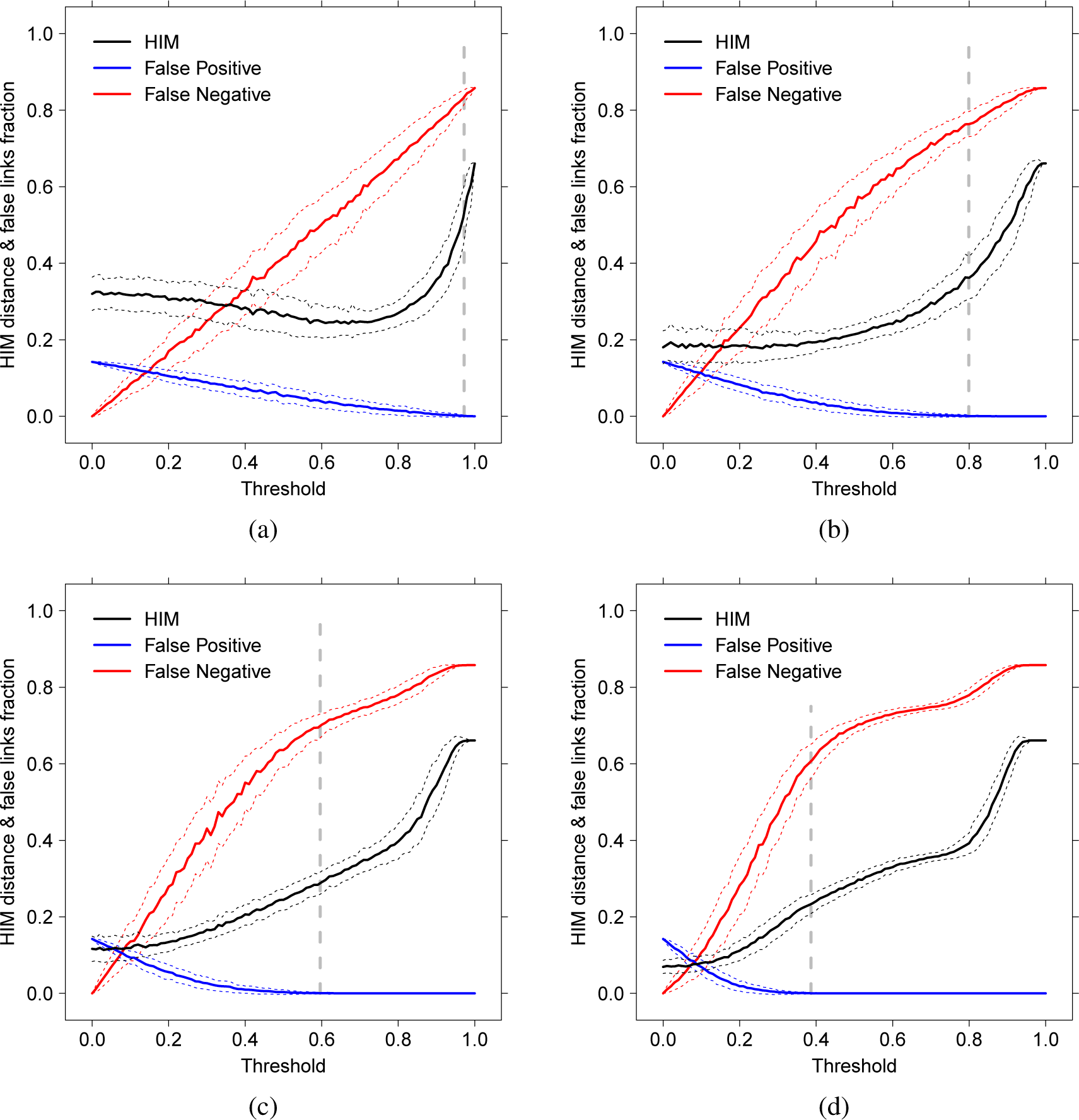
Coexpression inference of the coexpression network from subsampling of the *𝒪*_*T*_ dataset, on 5 (a), 10 (b), 20 (c) and 50 (d) samples. Solid lines indicate mean over 500 replicates of HIM distance (black), ratio of False Positive (blue) and ratio of False Negative (red); dotted lines of the same color indicate +/−1*σ*, while grey vertical dashed lines correspond to the secure threshold *p̄*.

Again, the secure threshold *p̄* corresponds to the smallest Pearson value warranting no false positive links included. Moreover, in almost all displayed situations, the threshold *p̄* is approximately the value where the HIM distance starts growing quicker, while the false positive rate remains under 0.8.

### 4.2 Small sample size

When the sample size is very small, the novel hard thresholding introduced here can severely limit the conclusions than can be drawn without incurring in the risk of discussing false positive links. This problem can be particularly evident in differential network analysis tasks [71, 72, 73, 31, 74], where loosening the threshold may lead to consider unsupported variations between networks in different conditions. In what follows we show two cases of (almost) negative results, where the experimental conditions tightly bound the possible differential coexpression network analysis.

#### Pancreatic Cancer

The first example is based on a pancreatic cancer dataset, publicly available at GEO http://www.ncbi.nlm.nih.gov/geo/, at the accession number GDS4329 and originally analysed in [58]. The dataset consists of 24 samples from 6 patients suffering from pancreatic ductal adenocarcinoma, divided in 4 subgroup samples, *i.e.*, circulating tumor cells (C), haematological cells (G), original tumour (T), and non-tumoural pancreatic control tissue (P). The aim of the original study was to develop a circulating tumor cells gene signature and to assess its prognostic relevance after surgery, while here we concentrate on the feasibility of a differential coexpression network analysis. Namely, we explore the Pearson correlation networks build separately on the four classes of samples on a specific set of genes *S*, defined by the differential expression analysis. In particular, the set *S* include as nodes the genes resulting upregulated in the C subgroup and associated with both the p38 mitogen-activated protein kinase (MAPK) signaling pathway and the cell motility pathway, which were ranked as the pathways with the highest expression ratio. In details, the nine genes are Talin-1 (TLN1), signal transducer and activator of transcription 3 (STAT3), Vinculin (VCL), CCL5, autocrine motility factor receptor (AMFR), Tropomyosin alpha-4 chain (TPM4), arachidonate 12-lipoxygenase (ALOX12), Rho-guanine nucleotide exchange factor 2 (ARHGEF2), and engulfment and cell motility protein (ELMO1), respectively denoted by 1, …, 9 in the plots.

Following the formula in Definition 5, the secure threshold for nine genes and six samples is 0.8508: hard thresholding the four coexpression networks results in the graphs collected in Fig. 8. As shown by the plots, the number of edges that result statistically significant over the secure threshold 0.8508 is small: namely 6 for the class G, 4 for the classes C and P and none for the primary tumoral cells T. In particular, the classes C and G share the links VCL–CCL5 and VCL–ALOX12, while P and G share the link TPM4–ALOX12 and P and C have no common links. Clearly, the paucity of statistically significant links prevents any further quantitative comparison: in Fig 9 we show, for each networks, the number of links at a given correlation.

**Figure 8:**
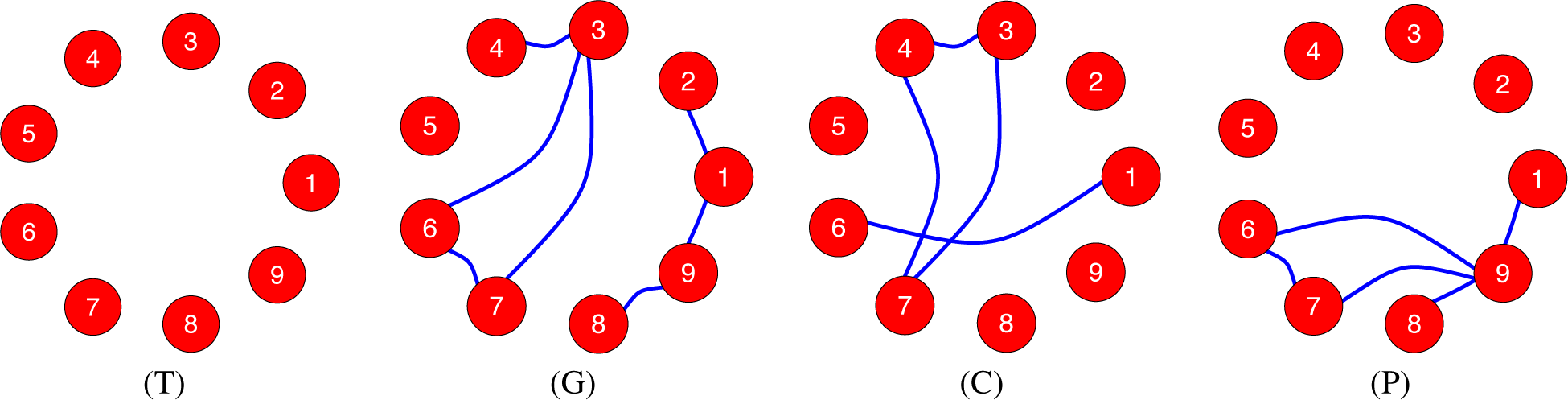
Correlation networks on the set *S* for the four classes T, G, C and P, thresholded at Pearson correlation coefficien 0.8508.

**Figure 9:**
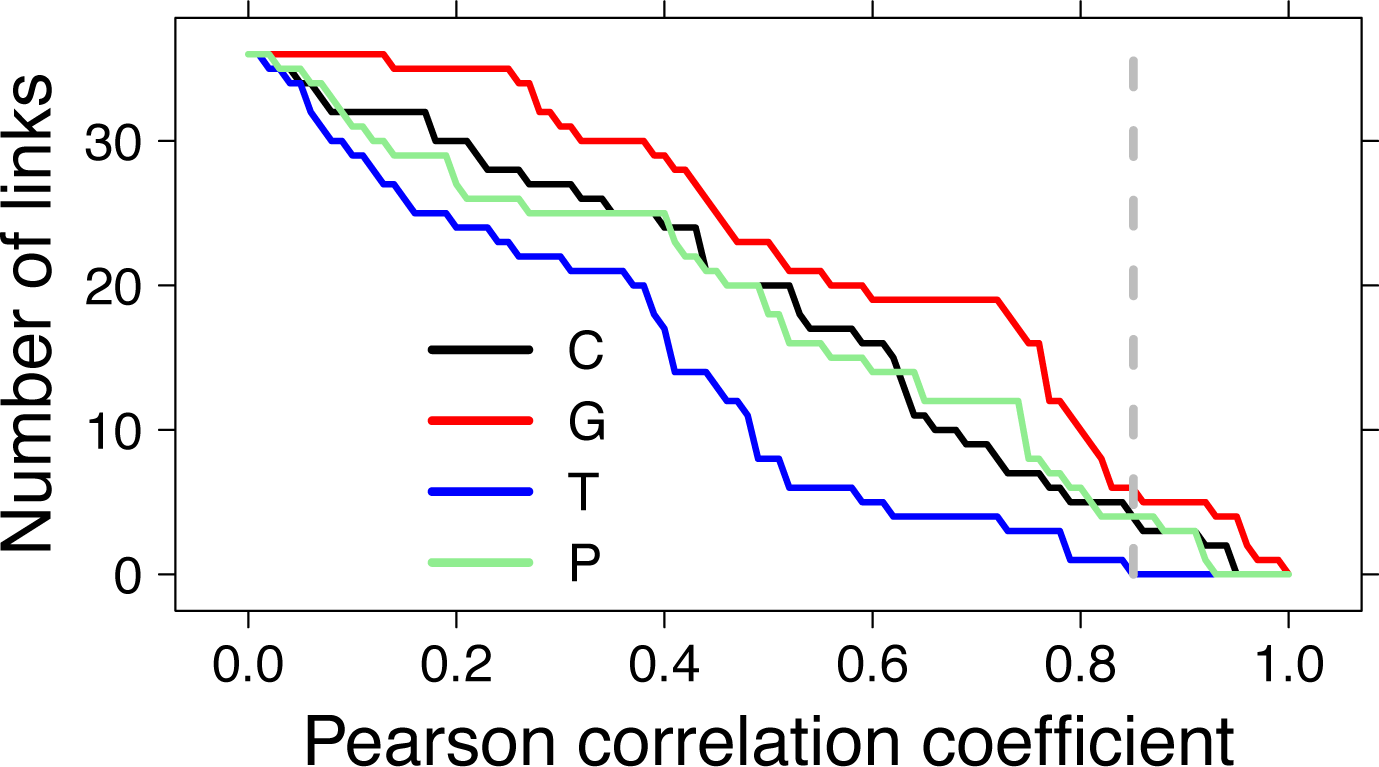
Number of links with correlation values larger than a given threshold for the coexpression networks C, P, T, and G; the vertical gray dashed line corresponds to Pearson correlation 0.8508, the secure threshold for 9 nodes and 6 samples.

#### Alzheimer data

A similar situation occurs with the Alzheimer dataset studied in [59, 60, 61] and available at GEO http://www.ncbi.nlm.nih.gov/geo/, at the accession number GSE4226. The dataset collect the expression of peripheral blood mononuclear cells from normal elderly control (NEC) and Alzheimer disease (AD) subjects. The NEC and AD subjects were matched for age and education; the Mini-Mental State Examination (MMSE) [75] was administered to all subjects, and the mean MMSE score of the AD group was significantly lower than that of the NEC subjects. Targets from biological replicates of female (F) and male (M) NEC and female and male AD were generated and the expression profiles were determined using the NIA Human MGC custom cDNA microarray. Each combinations of the sex and disease phenotypes has a cohort size of seven samples.

The original aim of the studies was the comparison between NEC and AD and the identification of genes with disease and gender expression patterns. In what follows, we show that, given the small sample size, very little can be assessed by a differential coexpression network analysis (see [76] for a recent larger miRNA coexpression study on a cohort of 363 individuals). In particular, from the KEGG Database http://www.genome.jp/kegg/ [77, 78] we extracted the Alzheimer’s disease pathway in Homo sapiens (KEGG accession hsa05010) and we extracted, from the original 32 genes included in the pathway, the 10 genes spotted on the platform with no missing value across the 28 total samples. The ten resulting genes are apolipoprotein E (APOE), amyloid beta (A4) precursor protein (APP), glycogen synthase kinase 3 beta (GSK3B), cyclin-dependent kinase 5 (CDK5), microtubule-associated protein tau (MAPT), presenilin 2 (Alzheimer disease 4) (PSEN2), amyloid beta A4 precursor protein-binding, family B, member 1 Fe65 (APBB1), lipoprotein lipase (LPL), synuclein alpha non A4 component of amyloid precursor (SNCA) and anterior pharynx defective 1 homolog A (APH1A), numbered from 1 to 10 in the above order in what follows. The resulting heatmap is shown in Fig. 10. The coexpression networks for the four combinations of sex (M/F) and disease (NEC/AD) are shown in Fig. 11, where the secure threshold is *p̄* = 0.8166. Again, the number of links whose correlation value is above the secure threshold is very small: however, all the retrieved links are well known in literature [79] and in dedicated webservers such as GeneMANIA http://www.genemania.org [80]. Clearly, if we consider the two main classes AD and NEC, the number of samples grows to 14 for each class, and the threshold *p̄* can be relaxed down to 0.5943. The two resulting networks are displayed in Fig. 12, together with the trend of the HIM distance between AD and NEC as a function of the threshold, both globally and separately for gender, where we can see that the selected threshold, in all cases, falls after the maximal distance between disease and control group. As a major effect emerging when comparing the coexpression network of the AD patients versus the NEC individuals we note that the connections between CDK5 and PSEN2, APBB1, LPL, SNCA are lost in the disease networks, while connections appear bewteen APP and APOE, CDK5, MAPT, PSEN2; changing of regualation of CDK5 and APP in AD patients are well known in literature: see for instance [81, 82, 83].

**Figure 10:**
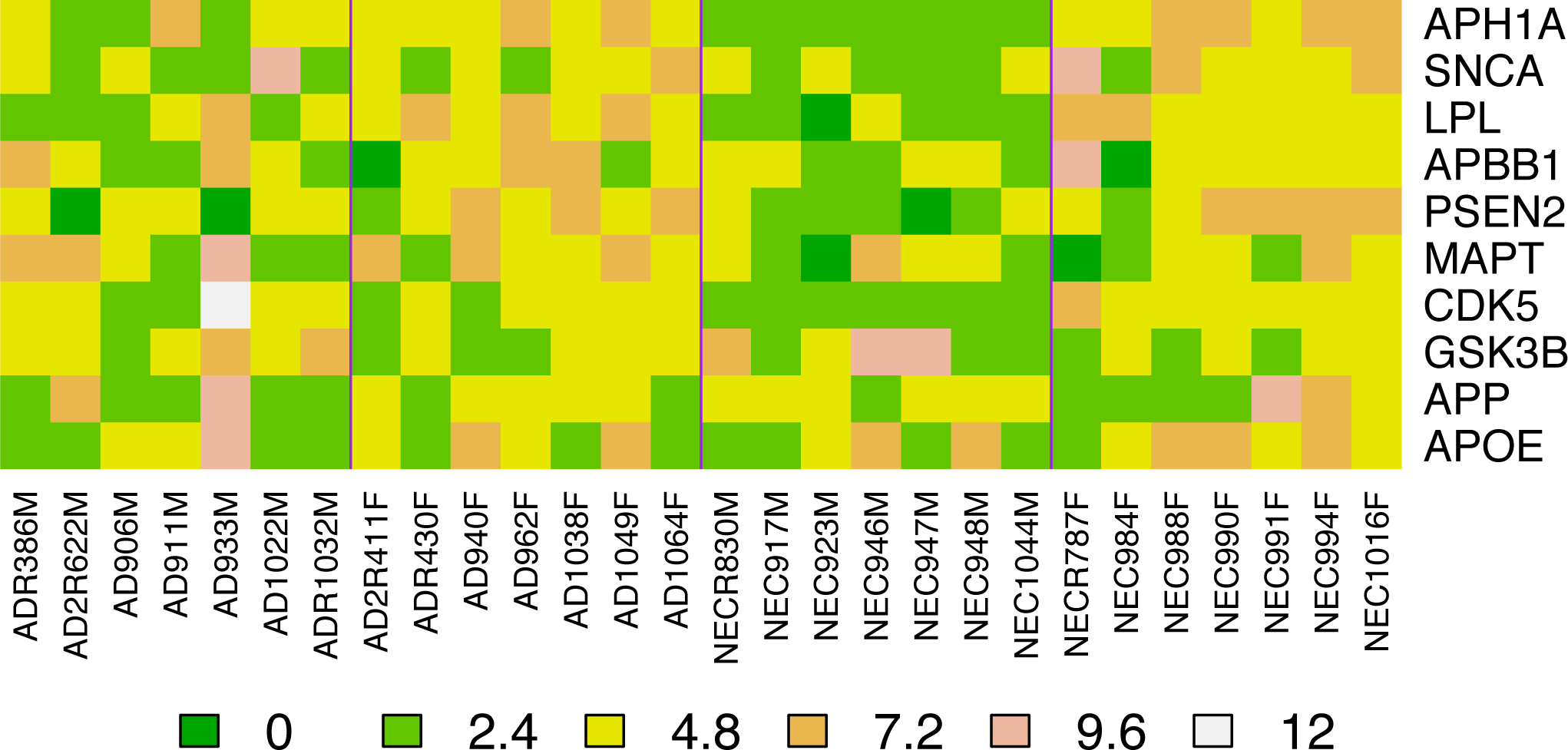
Heatmap of the expression of the ten genes of the Alzheimer pathway on the 28 samples of the Alzheimer dataset. Vertical lines separate samples groups.

**Figure 11:**
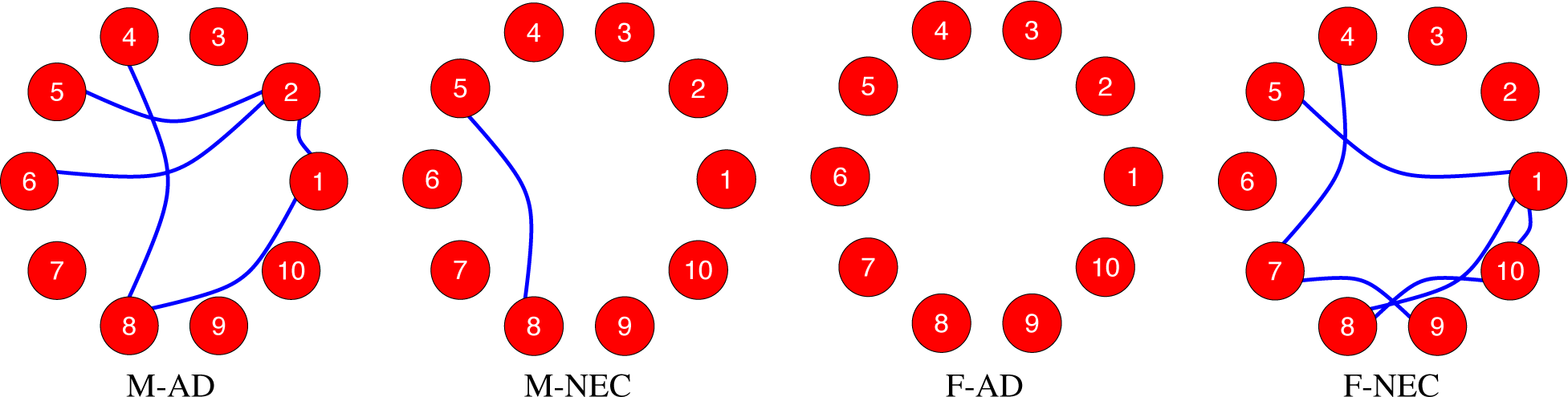
Correlation networks on the Alzheimer dataset *S* for the four classes M-AD, M-NEC, F-AD, F-NEC, thresholded at Pearson correlation coefficien 0.8166.

**Figure 12:**
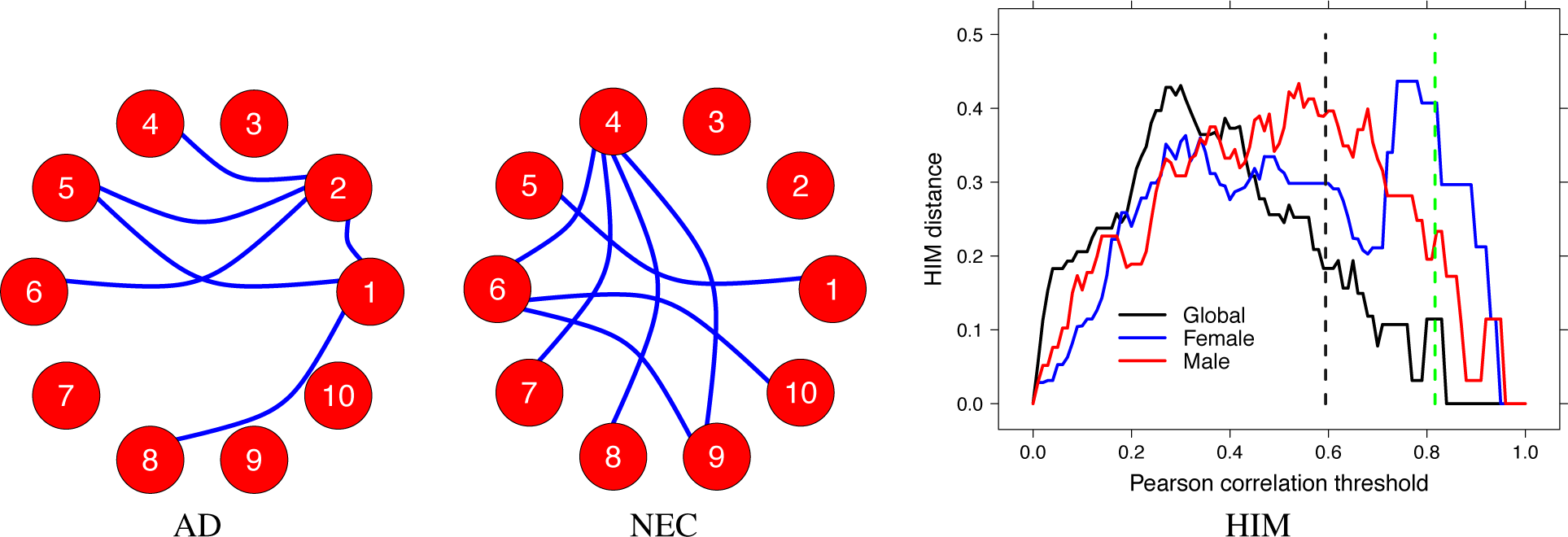
Alzheimer dataset: correlation networks for the two classes AD and NEC, thresholded at Pearson correlation coefficien 0.5943 (AD, NEC) and HIM curve of the distance between AD and NEC network versus the Pearson correlation threshold, globally (black) and separately for Male (red) and Female (blue) patients and controls. Grey dashed vertical line indicates the secure threshold *p̄* = 0.5943 for the global case, while the green line corresponds to the secure threshold *p̄* = 0.8166 for the sex disaggregated case.

## 5 Conclusion

A simple a priori, theoretical and non-parametric method is proposed for the selection of an hard threshold for the construction of correlation networks. This model is based on the requirements of filtering random data due to noise and reducing the number of false positive, and it is implemented by means of geometric properties of the Pearson correlation coefficient. This new approach can be especially useful in small sample size case, probably the most common situation in profiling studies in functional genomics. Finally, when the number of samples increase, coupling this method with soft thresholding approaches, can help recovering false negative links neglected by too strict thresholds.

## Footnotes

1 The HIM distance [69, 70] is a metric between networks having the same nodes, ranging between 0 for identical networks and 1, attained comparing the clique with the empty graph.

